# Detection and quantification of the histone code in the fungal genus *Aspergillus*

**DOI:** 10.1101/2023.01.31.526455

**Authors:** Xin Zhang, Roberta Noberini, Alessandro Vai, Tiziana Bonaldi, Michael F. Seidl, Jérȏme Collemare

## Abstract

In eukaryotes, the combination of different histone post-translational modifications (PTMs) – the histone code – impacts the chromatin organization as compact and transcriptionally silent heterochromatin or accessible and transcriptionally active euchromatin. Although specific histone PTMs have been studied in fungi, an overview of histone PTMs and their relative abundance is still lacking. Here, we used mass spectrometry to detect and quantify histone PTMs in germinating spores of three fungal species belonging to three distinct taxonomic sections of the genus *Aspergillus* (*Aspergillus niger, Aspergillus nidulans* (two strains), and *Aspergillus fumigatus*). We overall detected 23 different histone PTMs, including a majority of lysine methylations and acetylations, and 23 co-occurrence patterns of multiple histone PTMs. Among those, we report for the first time the detection of H3K79me1/2 and H4K5/8/12ac in Aspergilli. Although all three species harbour the same PTMs, we found significant differences in the relative abundance of H3K9me1/2/3, H3K14ac, H3K36me1 and H3K79me1, as well as the co-occurrence of acetylation on both K18 and K23 of histone H3 in a strain-specific manner. Our results provide novel insights about the underexplored complexity of the histone code in filamentous fungi, and its functional implications on genome architecture and gene regulation.

**Highlights:**

1. Quantitative mass spectrometry allows uncovering major histone PTMs.
2. H3K79me1/2 and H4K5/8/12ac are new histone PTMs detected in Aspergilli.
3. The relative abundance of histone PTMs varies in a strain-specific manner.
4. Variation in histone PTM abundance reflects different chromatin status.

## 1. Introduction

In eukaryotes, the genetic information is stored within the nucleus as chromatin, a complex of DNA and proteins of which the basic unit is the nucleosome (1). A nucleosome is composed of a histone octamer formed by two copies of each histone protein H2A, H2B, H3, and H4 wrapped by a DNA stretch of 145-147 bp (1). Nucleosomes are linked by 30-50 bp DNA sequence and the linker histone protein H1 (1). Histone proteins are highly conserved, consistent with their fundamental roles in structuring, scaffolding, and packaging DNA (2). Each histone protein contains a globular central domain that interacts with the nucleosomal DNA, and a lysine- and arginine-rich N-terminal tail that is subject to a diverse array of chemical modifications (2). These histone post-translational modifications (PTMs) contribute to the regulation of chromatin accessibility by two mechanisms: first, they directly alter chromatin packaging by changing histone proteins’ net charges; second, histone PTM-specific binding proteins are recruited and can influence inter-nucleosomal interactions (3). The resulting chromatin changes lead to transitions between inaccessible and transcriptionally silent chromatin regions, the so-called (facultative or constitutive) heterochromatin, and the accessible and transcriptionally active regions, the so-called euchromatin (3). The combination of different histone PTMs at a single genomic locus – the histone code – contributes to the accessibility of chromatin for the transcriptional machinery, thereby regulating gene expression to influence not only growth and development but also responses to environmental challenges (2).

In the last decades, a plethora of different histone PTMs have been uncovered in diverse organisms from plants to human, with an enrichment of histone mono-, di-, and tri-methylation (me) and acetylation (ac) (4). Accordingly, the vast majority of studies in fungi thus far have focused on these PTMs (5). Euchromatin has been typically associated with methylations on histone protein H3 at lysine 4 (K4), K36 and K79, and acetylations on H3 at K9, K14, K18, K23, and K27 and H4 at K5, K8, K12, and K16 (6,7). For instance, in the budding yeast *Saccharomyces cerevisiae* methylation on the lysine 4 of histone protein H3 (H3K4me) and H3K36me are enriched at the 5’ and 3’ end of actively transcribed genes, respectively, while H3K79me has been detected at the coding regions of active genes (8,9). H3K4ac locates just upstream of actively transcribed genes (10), and H3K9ac and H4K5ac have been shown to induce transcription elongation *in vitro* (11). In members of the filamentous Aspergilli, some of these euchromatin marks are shown to positively influence the production of secondary metabolites, including H3K4me in *Aspergillus oryzae* (12); H3K36me, H3K14/18/23ac (acetylation on either of these lysines), and H4K16ac in *Aspergillus flavus* (13,14); H3K9ac in *Aspergillus fumigatus* (15) and *Aspergillus nidulans* (16); and H3K27ac in *A. fumigatus* (17). Histone PTMs can also be linked to gene silencing, as first shown for H3K9me and H3K27me that are marks for constitutive heterochromatin in animals and plants (3). H3K9me3 plays a similar role in fungi as this PTM is typically located at telomeres, sub-telomeres, centromeres, and transposon-rich regions in the fission yeast *Schizosaccharomyces pombe* (18), the filamentous model fungus *Neurospora crassa* (19) and many other filamentous fungi (20). H3K9me also represses the transcription of secondary metabolite genes in *A. nidulans* (16) and *A. fumigatus* (21), and the deletion of the enzyme responsible for the deposition of this PTM leads to an increased level of secondary metabolite production. Facultative heterochromatin is linked with H3K27me3, and unsurprisingly, it does generally not overlap with the euchromatin mark H3K4me in the cereal pathogens *Fusarium graminearum* (22) and *Fusarium fujikuroi* (23). By contrast, SET7 (KMT6) that is responsible for H3K27me is absent in Aspergilli, and consequently, no H3K27me is detected (24). Although many PTMs have been clearly associated with gene activation or repression only, their effect can also be more subtle, depending on the genomic context. In *N. crassa* and *F. fujikuroi*, H3K36me catalyzed by SET-2 drives gene expression at the euchromatic region, while H3K36me catalyzed by ASH-1 at sub-telomeric regions is associated with gene repression (25,26). This complexity is even higher when considering the role of less abundant histone PTMs like histone acylation, phosphorylation, ubiquitination, and sumoylation, which thus far have been mostly reported in human (27). In fungi, such less abundant PTMs have been only reported scarcely in *S. cerevisiae* (diverse acylations, phosphorylation, ubiquitination, and sumoylation) and *A. nidulans* (phosphorylation, ubiquitination, and sumoylation), and the phosphorylation of histone H3 serine 10 correlates with chromosome condensation (28).

Large-scale mass spectrometry (MS) studies have been performed on bulk histones from model species such as human, yeast, and Arabidopsis to uncover the full pattern of histone PTMs and their interactions (29,30). However, studies in filamentous fungi have mostly focused on quantitative changes of targeted histone PTMs in mutant strains deficient in specific histone-modifying enzymes (31,32). For instance, mutants of histone H3 demethylase *KdmA* or *KdmB* in *A. nidulans* show increased H3K36me3 or H3K4me3, respectively (31,32). Our previous evolutionary analysis of 16 chromatin modifier complexes across the *Aspergillus* genus demonstrated the conservation of most of the catalytic genes responsible for a variety of histone PTMs (24). However, we still lack an overview of (co-)occurrence of histone PTMs in filamentous fungi, as well as a better understanding of the quantitative diversity of histone PTMs between different species. Here, we used quantitative MS to uncover histone PTMs in three *Aspergillus* species (*Aspergillus niger, A. nidulans*, and *A. fumigatus)* from distinct taxonomic sections and to report their estimated quantitative intra- and inter-species difference.

## 2. Materials and methods

### 2.1. Fungal strains and culture

*Aspergillus niger* (strain: NRRL3, CBS 120.49) and *A. fumigatus* (strain: Af293, CBS 126847) were grown on MEA (Malt Extract Agar; Malt Extract Agar Oxoid 50 g/L) medium at 30 °C and 25 °C, respectively, while *A. nidulans* (strains: wtpaba and TN02A25, kindly provided by Prof. Joseph Strauss) was grown on YAG (Yeast Agar Glucose; 5 g/L yeast extract, 15 g/L agar, 20 g/L D-glucose, 1 mL/L trace elements (Solution A: dissolve 10 g EDTA and 1 g FeSO_4_ × 7H_2_O in 80 mL water and adjust pH to 5.5; solution B: dissolve 4.4 g ZnSO_4_ × 7H_2_O, 1 g MnCl_2_ × 4H_2_O, 0.32 g CuSO_4_ ×6H_2_O, and 0.22 g (NH_4_)_6_Mo_7_O_24_ × 4H_2_O in 80mL water; combine solutions A and B, adjust pH to 6.5) medium at 37 °C.

Fungal spores were harvested from 4- or 5-day-old sporulating plates with a spreader in 12.5 mL ice-cold ACES (1.822 g/L N-(2-Acetamido)-2-aminoethanesulfonic acid, 0.2 mL/L Tween 80) buffer. The spore suspension was centrifuged at 2,500 xg at 4 °C for 5 minutes. The supernatant was discarded, and spores were resuspended in 25 mL ice-cold ACES buffer. The spore suspension was centrifuged at 2,500 xg at 4 °C for 5 minutes, and the supernatant was discarded. Spores were resuspended in 10 mL ice-cold ACES buffer, and spore concentration was determined with a hemocytometer (Marienfeld, 0.0025mm^2^) or spore counter, and adjusted to 10^7^ spores/mL. For the overnight culturing, 10^7^ to 10^8^ spores were inoculated in 50 mL liquid medium in a 250 mL flask and incubated at the same temperature as plate culture, 200 rpm, for 16 hours. Germinated spores were centrifugated at 4,000 xg for 10 minutes and the supernatant was discarded. The germinated spore pellet was washed with 25 mL 0.8 M NaCl, followed by centrifugation at 4,000 xg for 10 minutes to collect them for histone protein extraction.

### 2.3. Histone protein extraction

Histone proteins were extracted according to an adapted histone protein extraction protocol (33). Briefly, the fungal material was freeze-dried, ground with a mortar and pestle, and further disrupted by TissueLyser II (QIAGEN, C.0659). 100 mg ground sample was first resuspended in 1 mL lysis buffer (1×PBS, 0.5 mM PMSF, 5 μM Leupeptin, 5 μM Aprotinin, and 5mM Na-butyrate) with high (10%) Triton X-100 concentration and incubated on an orbital shaker at 4 °C for half an hour. Then 10 mL lysis buffer with low (1%) Triton X-100 concentration was added and the sample was incubated for another half an hour. The samples were subsequently sonicated for 30 seconds with 10 repeats and 30-second break in between each repeat. The crude histone proteins were dissolved in 0.4 N H_2_SO_4_ and then precipitated in four volumes of acetone. Protein concentration was measured with the Bradford Protein Assay (34).

### 2.3. LC-MS/MS analysis of histone PTMs

Samples were separated on a 17% SDS-PAGE gel and histone bands were excised and in-gel digested using a double derivatization protocol that involves propionylation of lysines and N-terminal derivatization with phenyl isocyanate (35). Chemical propionylation of lysines (which occurs on unmodified or mono-methylated residues) impairs trypsin cleavage, resulting in proteolytic cleavage at arginine residues only and the generation of histone peptides of proper length for MS analysis. Peptide mixtures were separated by reversed-phase liquid chromatography and analyzed by MS on an Orbitrap instrument, as previously described (24).

The acquired RAW data were analyzed using the integrated MaxQuant software v.1.6.10. The Uniprot UP000000560, UP000006706, and UP000002530 databases were used for the identification of *A. nidulans, A. niger*, and *A. fumigatus* histone peptides, respectively. Enzyme specificity was set to Arg-C. The estimated false discovery rate was set at a maximum of 1% at both peptide and protein levels. The mass tolerance was set to 4.5 ppm for precursor and fragment ions. Two missed cleavages were allowed, and the minimum peptide length was set to four amino acids. The second peptide search was enabled. Variable modifications include lysine propionylation (+56.026215 Da), monomethylation-propionylation (+70.041865 Da), dimethylation (+28.031300 Da), trimethylation (+42.046950 Da), acetylation (+42.010565 Da), malonylation (+86.000394 Da), and succinylation (+100.016044 Da). N-terminal PIC labeling (+119.037114 Da) was set as a fixed modification (35). To reduce the search time and the rate of false positives, which increases with a higher number of variable modifications included in the database search, the raw data were analyzed through multiple parallel MaxQuant jobs (36), setting different combinations of variable modifications. Peptides identified by MaxQuant with Andromeda score higher than 50 and localization probability score higher than 0.75 were quantitated, either manually or by using a version of the EpiProfile 2.0 software (37) adapted to the analysis of histones from *Aspergillus* strains. Identifications and retention times were used to guide the manual quantification of each modified peptide using QualBrowser version 2.0.7 (Thermo Scientific). Site assignment was evaluated from MS2 spectra using QualBrowser and MaxQuant Viewer. Extracted ion chromatograms were constructed for each doubly charged precursor, based on its *m/z* value, using a mass tolerance of 10 ppm. For each modified histone peptide, the relative abundance was estimated by dividing the area under the curve (AUC) of each modified peptide by the sum of the areas corresponding to all the observed forms of that peptide (38). The MS proteomics data have been deposited to the ProteomeXchange Consortium (39) *via* the PRIDE partner repository with the dataset identifier PXD033478.

PCA figures were generated by sklearn.decomposition.PCA of Sklearn (40) in Python, heatmap and point plots were made using Seaborn (41) in Python. ANOVA test was performed by Bioinfokit (42) in Python.

## 3. Results and discussion

The study of histone PTMs in filamentous fungi has mostly been restricted to targeted analyses based on knowledge from other organisms. However, the discovery of novel histone PTMs and their potential co-occurrence on a single nucleosome requires the use of untargeted strategies. We here applied a quantitative MS approach to identify the (co-)occurrence of histone PTMs and to quantify their relative abundance in three *Aspergillus* species: *A. niger, A. fumigatus*, and *A. nidulans*, which belong to distinct taxonomic sections (43). Protein sequences of histones H2A, H2B, H3, and H4 from human, yeast, and the three Aspergilli show very high levels of sequence conservation, especially in the core domains of all histone proteins that fold into the characteristic bulk octamer around which the DNA strand wraps, as well as in the N-terminal tails of H3 and H4 that protrude from the surface of the nucleosome and harbors most of histone PTMs (44) (Figure 1). The majority of well-described histone PTMs, e.g., histone methylations and acetylations, occurs mostly on lysine residues, while other histone PTMs, such as phosphorylation, crotonylation, and ubiquitination, can also occur on other residues such as serine and threonine (30). For histone protein H3, we observed the conservation of all lysine residues in human, yeast, and Aspergilli, except for two additional lysine residues at positions 121 and 125 in *S. cerevisiae* (Figure 1). On these two residues, Cul8/Rtt101, an enzyme that is thought to be specific to *S. cerevisiae* as no homolog has yet been found in human or other fungi, can introduce histone ubiquitylation and thereby regulate nucleosome assembly (45,46). We similarly observed limited variation in the N-terminal tail of histone protein H4 and the global core domains of H2A and H2B, and most of the polymorphism occurs on amino acid residues that are not known to carry any histone PTM.

**Figure 1.**
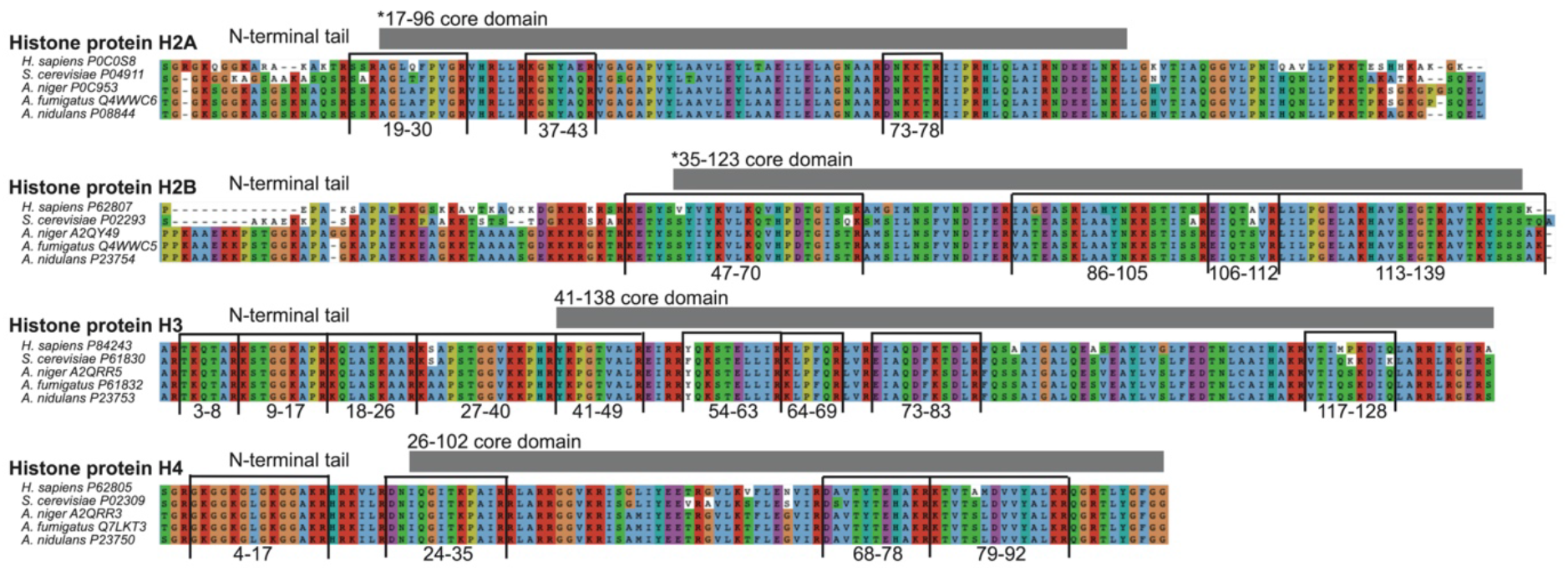
Histone proteins are well conserved in Aspergilli. Sequence alignment of histone proteins from human (H2A: P0C0S8, H2B: P62807, H3: P84243, and H4: P62805), yeast (*S. cerevisiae*) (H2A: P04911, H2B: P02293, H3: P61830, and H4: P02309), and *A. niger* (H2A: P0C953, H2B: A2QY49, H3: A2QRR5, and H4: A2QRR3), *A. fumigatus* (H2A: Q4WWC6, H2B: Q4WWC5, H3: P61832, and H4: Q7LKT3) and *A. nidulans* (H2A: P08844, H2B: P23754, H3: P23753, and H4: P23750, retrieved from UniProt database. The highly conserved globular histone core regions are highlighted by grey boxes with numbers indicating their position in human histone proteins with the symbol * showing the mismatch of the positions between human and Aspergilli histone proteins H2A and H2B caused by the poor alignment of the N-terminal tails. Peptides detected by MS in this study are labeled by black lines, and their relative positions in Aspergilli histone proteins are indicated (Table S1).

### 3.1. Histone protein extraction and digestion for MS-based analyses

We adapted a well-established protocol for human primary cells (47) to specifically extract histone proteins from filamentous fungi with three additional steps (freeze drying, grinding, and homogenization) for efficient disruption of fungal cell walls and release of nuclei (24,33) (Figure 2A). We extracted histone proteins in three biological replicates from the three *Aspergillus* species; two strains of *A. nidulans* (Wtpaba and TN02A25) were cultured on YAG medium, while *A. niger* strain NRRL3 and *A. fumigatus* strain Af293 were grown on MEA medium (Figure 2B). After visualizing the extracted histone protein on SDS-PAGE gel, bands ranging from 10,000 to 16,000 Dalton, which correspond to the size of histone proteins (48), were cut for in-gel digestion (Figure 2C). We used lysine derivatization with propionic anhydride on unmodified and mono-methylated lysines, followed by trypsin digestion and peptide N-terminal derivatization with phenyl isocyanate (PIC) to obtain histone peptides of amino acids length required for MS analysis. This method allows performing an ‘Arg-C-like’ digestion of histone proteins directly in the gel, which is the ideal digestion pattern for histone PTM analyses (35). In addition, because the derivatization occurs only on unmodified or monomethylated lysines, but not on di- or tri-methylated residues, it allows better discrimination of isobaric peptides (35). The N-terminal derivatization with PIC also aids in the detection of small and hydrophilic peptides, as well as the chromatographic separation of differentially acetylated peptides, as described previously (35,49). After digestion of isolated histone proteins from Aspergilli, we detected 20 different peptides from the four core histone proteins and three peptides from the histone protein variant H2A.Z (Figure 1D, Table S1). These peptides cover most of the lysine residues that are known to harbor histone PTMs.

**Figure 2.**
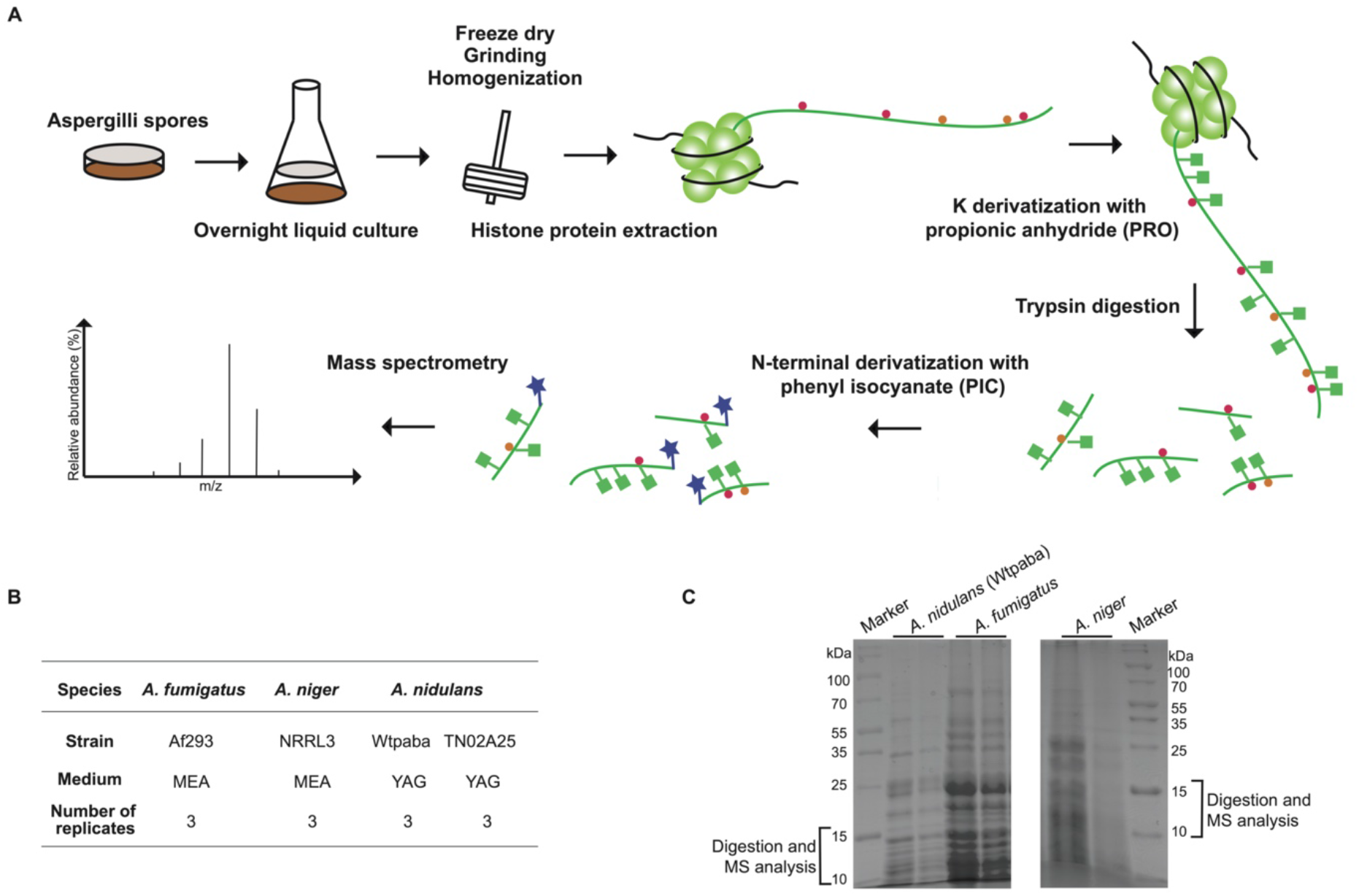
Histone proteins extraction for MS-based analysis of histone PTMs in Aspergilli. **A**. Schematic diagram of the experimental setup and workflow applied in this work. **B**. Summary of *Aspergillus* strains’ genetic background and culture conditions. The number of biological replicates for each condition is indicated. **C**. Histone proteins were extracted from the different *Aspergillus* species, and samples were visualized on the 17% SDS-PAGE gel. The region ranging from 10,000 to 16,000 Dalton was cut for the in-gel digestion step.

### 3.2. MS uncovers histone PTMs in three *Aspergillus* species

MS analyses resulted in the detection of 23 individual histone PTMs, as well as 23 distinct co-occurrence patterns of multiple histone PTMs that occur on the same peptide (Figure 3, Table S1). We did not detect any histone PTM on histone variant H2A.Z and histone H2A, but we were able to detect acetylation on K92 and co-occurrent mono-methylations on K98 and K99 on histone protein H2B (Table S1). These PTMs correspond to histone PTMs H2BR79ac (R: arginine) and H2BK85me1R86me1 in human, yet the functions of these PTMs are unknown (50). For histone PTMs on histone proteins H3 and H4 in Aspergilli, we successfully detected methylations on histone H3 at lysines 4, 9, 36, and 79, acetylation on histone H3 at lysines 9, 14, 18, 23, 27, and 56, and histone H4 at lysines 5, 8, 12, 16, and 31 (Figure 3). All these modifications have previously been reported in human (38,51), Arabidopsis (52), *S. cerevisiae* (53), and in a few filamentous fungi (5,20,54). However, it is to our knowledge the first time that H3K79me1, H3K79me2, H4K5ac, H4K8ac, H4K12ac, and H4K31ac are detected in Aspergilli (Figure 3). Consistently with the occurrence of these PTMs, the catalytic enzymes that are responsible to write these PTMs are also present in Aspergilli: Dot1 (Disruptor of telomeric silencing 1) methylates H3K79 and NuA4 (Nucleosomal Acetyltransferase of histone H4)/NuB4 (Nuclear Hat1p-containing type B histone acetyltransferase) complexes acetylate H4K5, H4K12, and H4K16 (24). The Elongator complex and Sas2 enzyme that are present in Aspergilli genomes are likely responsible for H4K8ac and H4K16ac, respectively (24). In the human parasite *Toxoplasma*, H4K31ac is catalyzed by the acetyltransferase Gcn5 (55). As this enzyme is conserved in Aspergilli and involved in the acetylation of H3 lysine residues, it is likely that Gcn5 also catalyzes the acetylation of H4K31 in Aspergilli.

**Figure 3.**
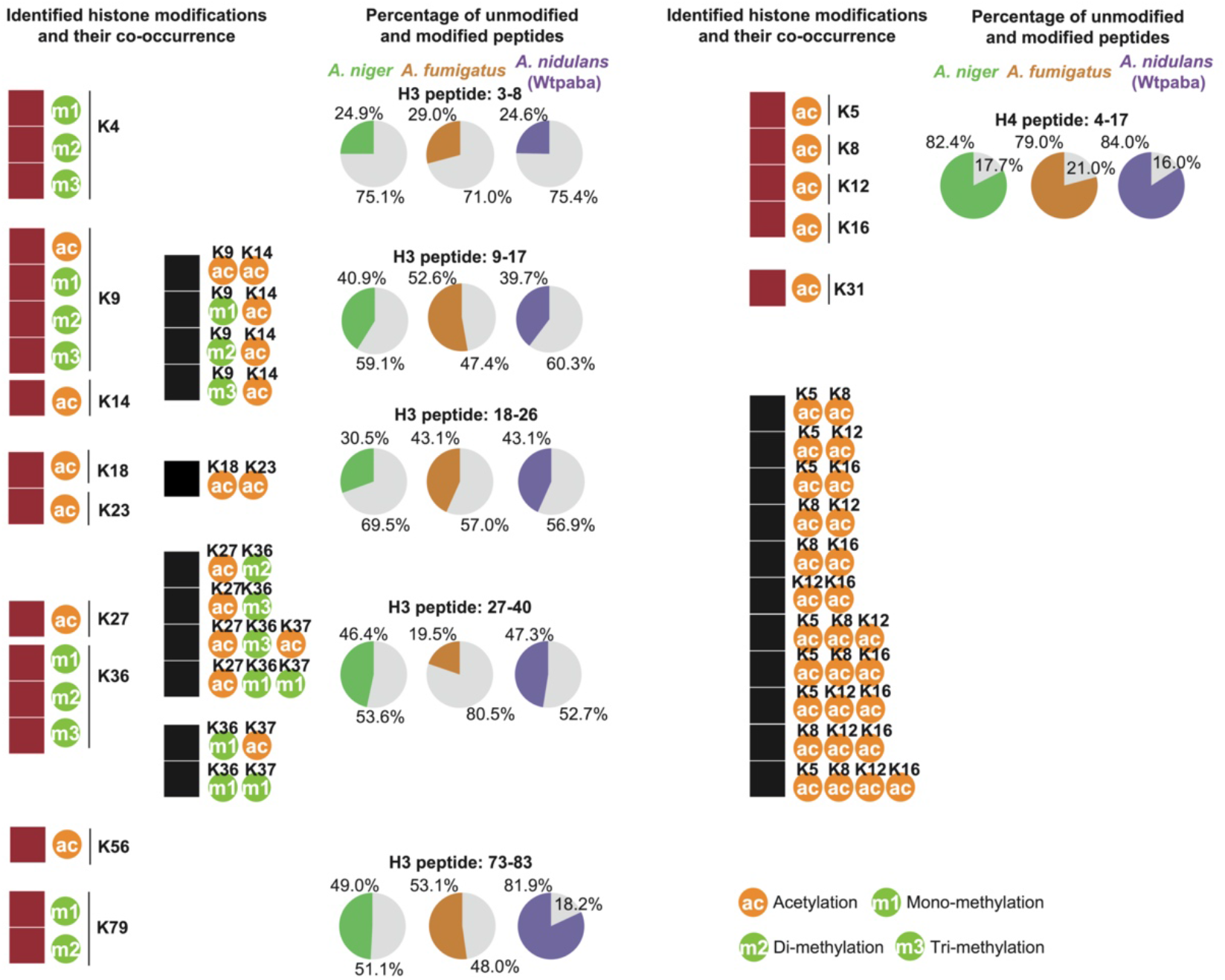
Identification of single and co-occurrent histone PTMs in Aspergilli. The red boxes show the identification of histone mono-, di-, and tri-methylation and acetylation while the black boxes show their co-occurrences in Aspergilli. Pie charts show the average relative abundance of unmodified peptides compared with modified peptides for each *Aspergillus* species. Unmodified peptides are shown in gray, and green, orange, as well as purple representing the proportion of modified peptides in *A. niger, A. fumigatus*, and *A. nidulans* (strain Wtpaba), respectively.

While we detected most of the known PTMs as well as a few additional ones for the first time in Aspergilli, we did not detect some histone PTMs which were expected to occur. Several reasons can explain these observations. First, PTMs might not be identified because they are under the detection threshold used in both the instrument and analysis pipeline. This reason likely explains why we did not detect any low-abundant histone PTMs. The abundance of certain expected PTMs like H3K79me3, H3K4ac, and H3K36ac could be higher and detectable under different growth conditions or developmental stages. Second, these PTMs are true absences in Aspergilli. We were not able to detect any methylation on H3K27, H4K5, H4K8, and H4K12 (Figure S1), which is expected due to the absence in Aspergilli of their catalytic enzymes, PRC2 (20,24) and SET5 (24) (Figure S2), respectively. Third, the mutual exclusivity of different histone PTMs on the same residue could lead to biased occupancy by only one histone PTM. For instance, apart from H3K9, either acetylation or methylation was detected on H3 lysines (Figure 3 and Figure S1). Such an explanation would indicate very homogenous samples regarding their chromatin status and PTMs. Last, antagonistic relationships between histone PTMs might contribute to some absences. For example, H4K16ac prevents H4K20me (56), explaining why we could detect the former but not the latter histone PTM in Aspergilli.

We also detected the co-occurrence of multiple histone PTMs on the same peptide (Figure 3). For example, we observed the co-occurrence of acetylations on different residues in the same peptide such as H3K9acK14ac, H3K18acK23ac, and H4 lysines 5, 8, 12, and 16 harboring acetylations at the same time on two, three, or four residues (Figure 3). Acetylation can also co-occur with methylation, such as H3K9me1/2/3K14ac, H3K27acK36me2/3, and H3K36me1K37ac, while H3K36me1 can co-occur with another mono-methylation on H3K37 (Figure 3). The detection and functional characterization of some of these co-occurrent patterns have been performed in several model species. For instance, in mouse, H3K9meK14ac has a role in gene silencing (57). Accordingly, double knockout of enzymes for H3K9acK14ac leads to the up-regulation of penicillin, sterigmatocystin, and orsellinic acid in *A. nidulans* (58). H3K18acK23ac was first reported to be acetylated by SAGA/ADA complex in *S. cerevisiae* (59) and then detected in human (60). The investigation in *S. cerevisiae* of the combinatorial effect of acetylation on H4 lysines 5, 8, 12, and 16 showed that they are redundant and function together as a group to induce gene activation instead of forming distinct combinational patterns (61). Even though the co-occurrence of histone PTMs has not yet been systematically studied, our data as well as the results from other model systems, such as human or mouse, provide valuable clues for the possible functions of co-occurring histone PTMs in gene regulation in Aspergilli.

Although relative abundances only represent an indication of the absolute abundance of a modification (since peptide detection by MS is affected by the peptide physicochemical properties and the digestion strategy (PMID: 25000943)), our data shows that the majority of detected histone H3 peptides is unmodified, while histone protein H4 exhibits a high percentage (79%-84%) of acetylated peptides in all three Aspergilli (Figure 3). The pattern is similar among the three species, with only a few observable differences. Fewer H3 peptides 18-26 are modified in *A. niger* (30% vs 43% in both *A. nidulans* and *A. fumigatus*), suggesting a lower abundance of H3K18ac and/or H3K23ac in this species. In contrast, *A. nidulans* exhibit more modified H3 peptides 73-83 (82% vs 49-53% in the other species), suggesting an enrichment in H3K79 methylation in this species. *A. fumigatus* differs from the two other species by having more modified H3 peptides 9-17 (53% vs 40-41%) and fewer modified H3 peptides 27-40 (19% vs 46-47%) (Figure 3). These differences suggest a higher abundance of H3K9 and/or H3K14 PTMs and a lower abundance of H3K27, H3K36, and/or H3K37 PTMS in *A. fumigatus*. These results collectively suggest that, although the same histone PTMs are detected in all three species, their abundances likely differ.

### 3.3. *Aspergillus* species differ in the relative abundance of histone PTMs

To uncover quantitative differences between histone PTMs, we calculated their relative abundance in three *Aspergillus* species (*A. niger, A. fumigatus*, and *A. nidulans* (wtpaba strain)) and plotted the figure of Principal Component Analysis (PCA) (Figure 4A). We observed that *Aspergillus* species are well separated in the PCA, suggesting that the three Aspergilli indeed differ quantitatively in their histone PTMs. These differences could be caused by differences in growth conditions (Figure 2B). However, we observed that even though *A. fumigatus* and *A. niger* were grown on the same medium compared with *A. nidulans, A. fumigatus* was more distinct from *A. nidulans* and *A. niger*. These distinctions suggest that the observed differences are not related to the different media but seem to be consistent with their taxonomic relationships. Specifically, *A. fumigatus* harbors significantly more H3K9 methylations, either alone or co-occurring with H3K14ac, and it has significantly less H3K36me1 compared to the other two Aspergilli (Figure 4B, Table S3). Although not significant but consistent with the observed proportions of modified peptides, *A. fumigatus* appears enriched in H3K79me1 (Figure 3 and 4B). In contrast, H3K9me1/2K14ac showed lower abundance in *A. niger* compared to the other two species, and H4K8ac was significantly higher in *A. fumigatus* compared to *A. nidulans*. Although non-significant but expected from the proportion of modified peptides, *A. niger* tends to contain fewer H3K18ac and H3K23ac and *A. nidulans* appears particularly rich in H3K79me2 and poor in H3K79me1 (Figure 3 and 4B). The observed differences suggest that *A. fumigatus* exhibit more heterochromatin-linked modifications compared to the other two species as it harbors more H3K9me1/2/3 linked with constitutive heterochromatin (62) and H3K9meK14ac associated with triggering gene silencing (57). For euchromatin, *A. fumigatus* uses more H3K79me1 and the combination of acetylations on diverse lysines of histone protein H4, while *A. niger* and *A. nidulans* harbors more H3K36me1 (Figure 4B). Even though we observed no qualitative difference in histone PTMs among three Aspergilli, these species differ quantitatively in specific histone PTM abundances, suggesting that the three populations of germinating spores (and their replicates) were very homogenous and in a similar state and they used a mix of marks associated with gene activation and gene silencing. Yet, the absence or low abundance of histone PTMs like H3K4ac, H3K14me, or H3K36ac could indicate their important roles in gene regulation in other developmental stages.

**Figure 4.**
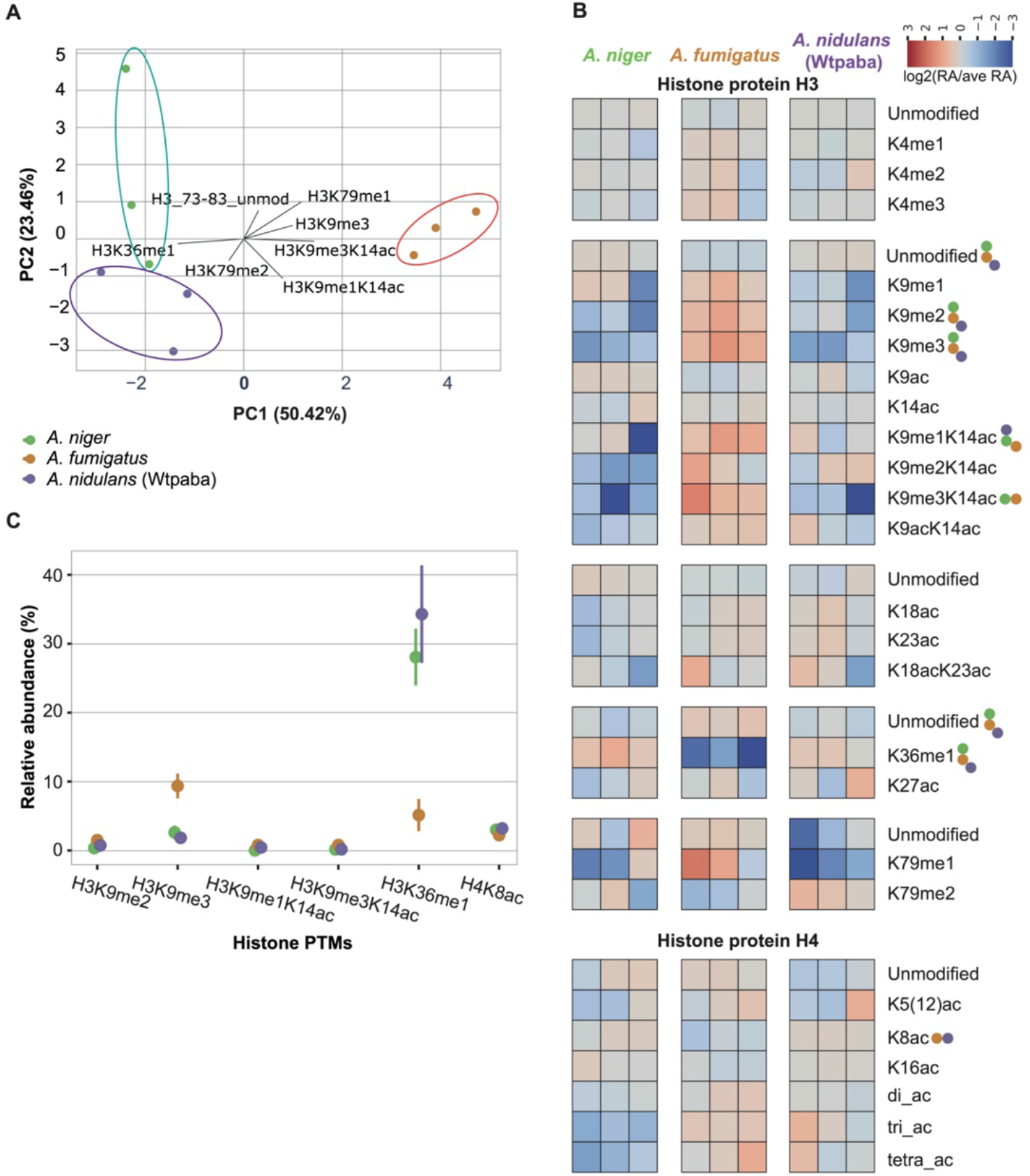
Quantification of histone PTMs in three *Aspergillus* species. **A**. Principal component analysis (PCA) of three Aspergilli based on their relative abundance of histone PTMs (Table S3). **B**. Quantitative comparison of all histone PTMs detected by MS. The heatmap displays red or blue color boxes according to the log_2_(relative abundance/average of relative abundance) of different peptides in Aspergilli. ‘Di-ac’ indicates H4K12acK16ac, H4K5acK12ac, H4K5acK16ac, H4K5acK8ac, or H4K8acK16ac; ‘tri-ac’ means H4K5acK12acK16ac, H4K5acK8ac12ac, H4K5acK8acK16ac, or H4K8acK12acK16ac; and ‘tetra-ac’ indicates H4K5acK8ack12acK16ac. To determine significant differences between the three *Aspergilli* species, we applied ANOVA tests; p-values < 0.025 were deemed significant (three biological replicated per species); colored dots indicate significant differences between represented species. **C**. Relative abundance of histone PTMs that are significantly different. Points show the average relative abundance and lines indicate standard deviation.

### 3.4. *A. nidulans* strains differ quantitatively in euchromatin marks

To evaluate the intra-species variation in the relative abundance of histone PTMs, we compared the MS data of another *A. nidulans* strain (TN02A25) to Wtpaba (Figure 5). In general, we observed that TN02A25 spurs the same type of histone PTMs as we observed in the other strains (Figure 4B and 5C), indicating all Aspergilli tested in this study used the same histone code. However, the two strains show a few differences, already when only considering the percentage of unmodified versus modified peptides. The vast majority of detected histone H3 peptides were unmodified (> 50%) and histone H4 peptides were highly acetylated in TN02A25 like the other strains (Figure 4 and Figure 5A). However, TN02A25 differs from Wtpaba in H3 peptides 27-40 (11% vs 47% modified peptides in Wtpaba) and 73-83 (46% vs 82% modified peptides in Wtpaba). These proportions in TN02A25 are similar to those found in *A. fumigatus* or *A. niger* and *A. fumigatus*, respectively (Figure 3), and suggest that the two *A. nidulans* strains differ in histone PTM abundance on those peptides.

**Figure 5.**
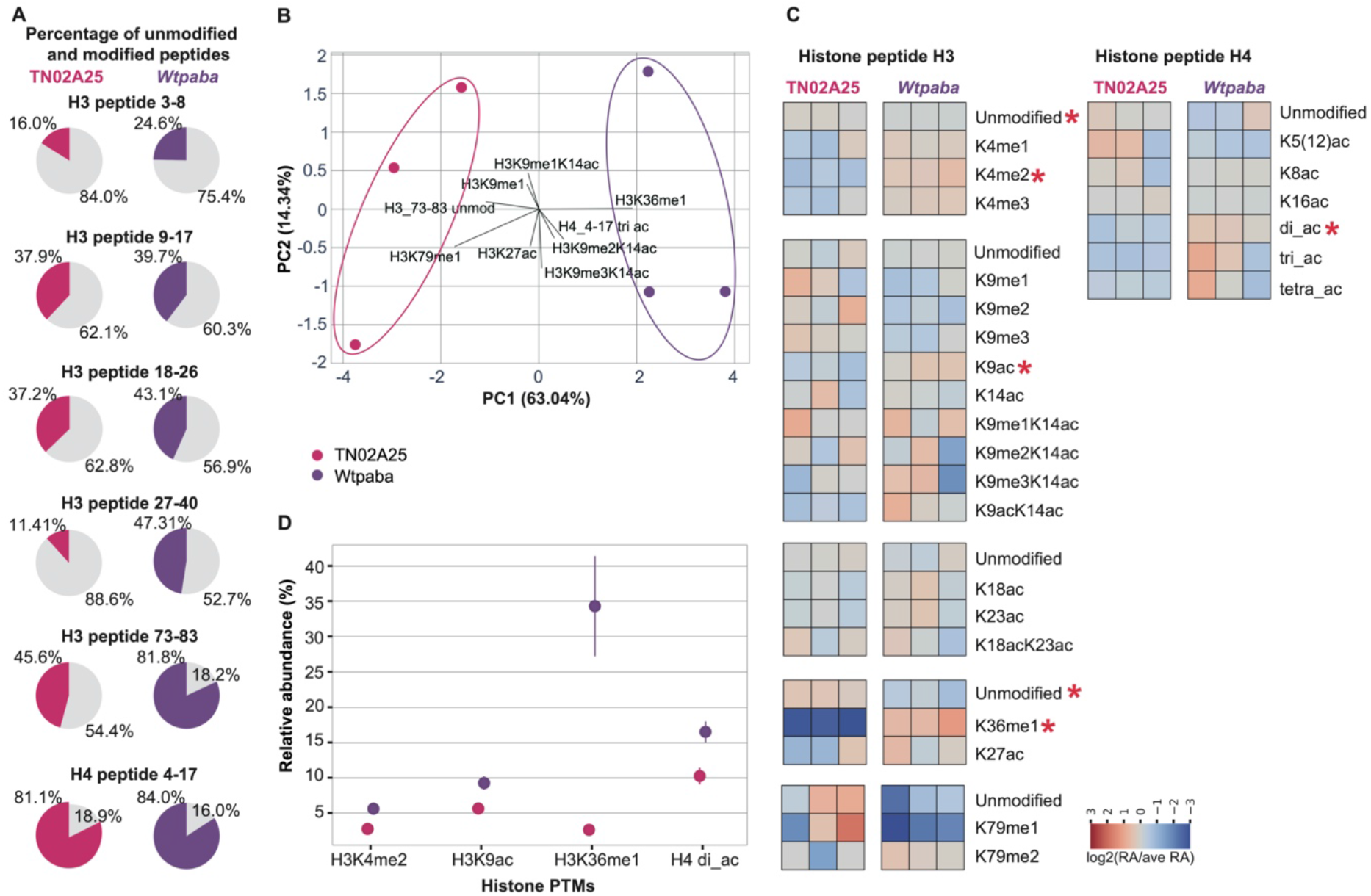
Quantification of histone modifications in two *A. nidulans* strains. **A**. The average relative abundance of unmodified peptides compared with modified peptides. Grey represents the unmodified peptide percentage in the pie chart, while pink and purple represent the modified peptide percentage of TN02A25 and Wtpaba *A. nidulans* strains, respectively. **B**. Principal component analysis (PCA) of two strains based on their relative abundance of histone PTMs. **C**. Quantitative comparison of all histone PTMs detected by MS. The heatmap displays red or blue color boxes according to the log_2_(relative abundance/average of relative abundance) of different peptides in two *A. nidulans* strains. ‘Di-ac’ indicates H4K12acK16ac, H4K5acK12ac, H4K5acK16ac, H4K5acK8ac, or H4K8acK16ac; ‘tri-ac’ means H4K5acK12acK16ac, H4K5acK8ac12ac, H4K5acK8acK16ac, or H4K8acK12acK16ac; and ‘tetra-ac’ indicates H4K5acK8ack12acK16ac. ANOVA test was performed to show the significance of the difference of one histone modification between two strains when p-value < 0.025. Red stars indicate significant differences. **D**. The top four histone modifications that are significantly different. Points are average relative abundance and lines are standard deviation.

The quantitative comparison of histone PTMs between the two *A. nidulans* strains shows good separation in the PCA analysis (Figure 5B). Consistent with previous results and analyses, TN02A25 exhibit significantly less H3K36me1 compared to the Wtpaba strain. Although not significant, the enrichment in H3K79me2 and lower abundance in H3K79me1 in Wtpaba contributes to the difference between both strains (Figure 5). In addition, H3K4me2, H3K9ac, H3K36me1, and H4di_ac exhibited significantly higher abundance in Wtpaba than TN02A25 (Figure 5C-D, Table S4). As these histone PTMs are all related to regulation of euchromatin.

Moreover, we observed the intra-species variability can be as high as the inter-species variability as the histone PTM abundances of Wtpaba and TN02A25 are more related to *A. niger* and *A. fumigatus*, respectively (Figure S3A). Four single or co-occurrent histone PTMs, H3K36me1, H3K79me1, H3K9me3K14ac, and H3K9me1K14ac displayed the most distinct abundant patterns among the four strains, followed by H3K9me1, H3K9me2, and H3K18acK23ac being less abundant in *A. niger* compared to the other three strains (Figure S3B). These marks showing distinct relative abundance agreed with the marks we pinpointed from both species and strain comparisons. Thus, the observed differences between species are likely a ‘strain effect’ and highlight the importance of including several strains in the study of histone PTMs.

The use of MS spectrometry is powerful to detect and quantify histone PTMs in filamentous fungi and such an approach is needed to understand the chromatin status at different developmental stages. The present study focused on germinating spores, a particular stage that requires the expression of a specific set of genes and is expected to not favor the production of secondary metabolites. Similar future studies in Aspergilli at different developmental stages and different culture conditions will fully uncover quantitative variations in histone PTMs and might lead to identifying key PTMs for certain applications, such as the controlled activation of secondary metabolite production. Finally, while this work has revealed quantitative differences between species and strains, it remains to determine whether they are due to different PTM abundance at the same loci, or whether they reflect a different number of loci covered by these PTMs, which would reflect activation or repression of different sets of genes. These future studies will contribute to understanding how single and co-occurring histone PTMs regulate gene expression in Aspergilli and more broadly in other filamentous fungi.

## Author contributions

X.Z. formal analysis, investigation, methodology, visualization, writing original draft; R.N. and A.V. formal analysis, investigation, methodology; T.B. funding acquisition, supervision, resources; J.C. conceptualization, funding acquisition, methodology, project administrations, supervision, visualization, writing original draft; M.F.S. conceptualization, funding acquisition, methodology, project administrations, supervision, visualization, writing original draft.

## Acknowledgments

We thank Jos Houbraken for providing *A. fumigatus* strain and Joseph Strauss for providing two *A. nidulans* strains.

## Appendix A. Supplementary materials

Supplementary Table 1 Histone peptides identified via mass spectrometry

Supplementary Table 2 Mass spectrometry result for Aspergilli

Supplementary Table 3 ANOVA test for all modifications in three Aspergilli

Supplementary Table 4 ANOVA test for all modifications in two *Aspergillus nidulans* strains

Supplementary Table 5 ANOVA test for all modifications in four *Aspergillus* strains

Supplementary Figure 1 Comparison of experimentally detected histone post-translational modifications (PTMs) in human, yeast, and Aspergilli

Supplementary Figure 2 SET5 was lost in Aspergilli

Supplementary Figure 3 Distinct abundant patterns of several histone PTMs differ different *Aspergillus* species/strains.

## Data availability

Data statement: All supporting data, code, and protocols have been provided within the article or through supplementary data files. Supplementary material is available with the online version of this article. The MS results are deposited in the PRIDE database with the dataset identifier PXD033478.

