## Supplementary material for "Detection and quantification of the histone code in the fungal genus *Aspergillus*": Figure S1

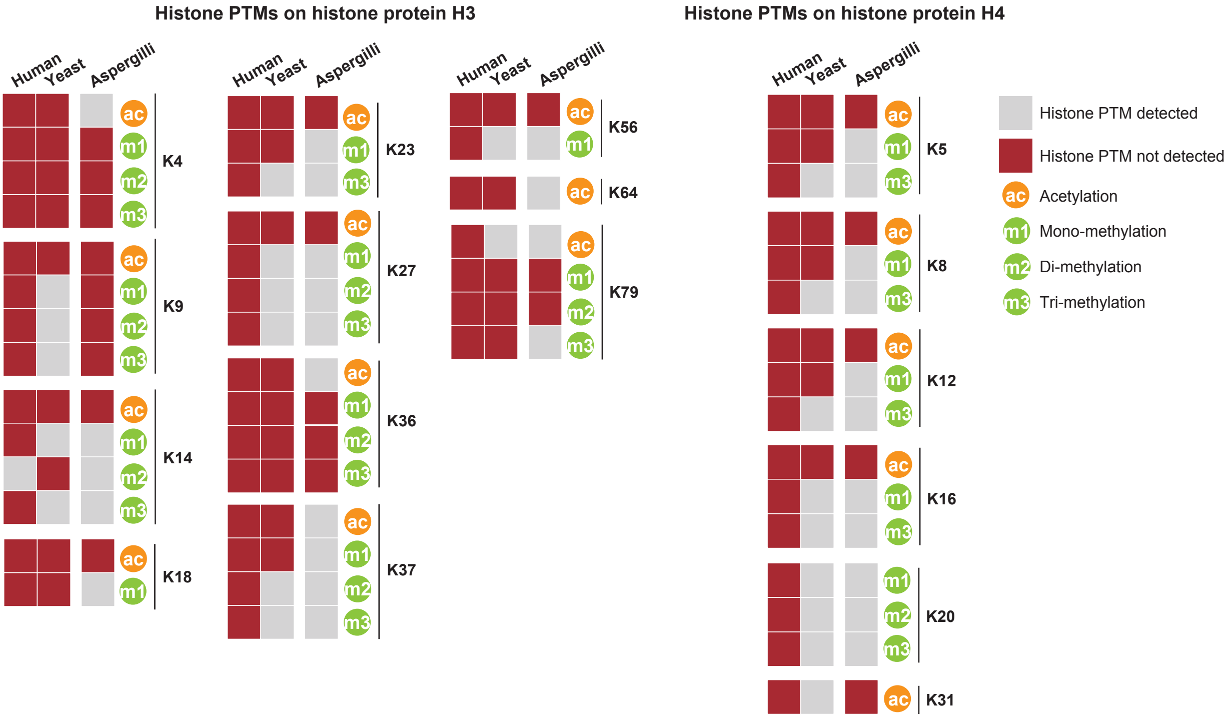


**Supplementary Figure 1 Comparison of experimentally detected histone post-translational modifications (PTMs) in human, yeast, and Aspergilli.** The summary of histone PTMS in human and yeast is adapted from a recently published paper (Grau-Bové, Xavier, et al., Nature Ecology & Evolution, 2022) with a few corrections: 1) H3K64ac can be detected in human (Pradeepa, Madapura M., et al., Nature genetics, 2016); 2) H3K9me1 is lost in yeast (Zhang, Xing, et al., Cell, 2002 and Marina, Diana B., et al., Genes & development, 2013); 3) H4K5me1 (Green, Erin M., et al., Nature structural & molecular biology, 2012) and 4) H4K8ac (Magraner-Pardo, Lorena, et al., BMC genomics, 2014) can be detected in yeast. The detection of histone PTMs in Aspergilli comes from this study.
