## Supplementary material for "Detection and quantification of the histone code in the fungal genus *Aspergillus*": Figure S2

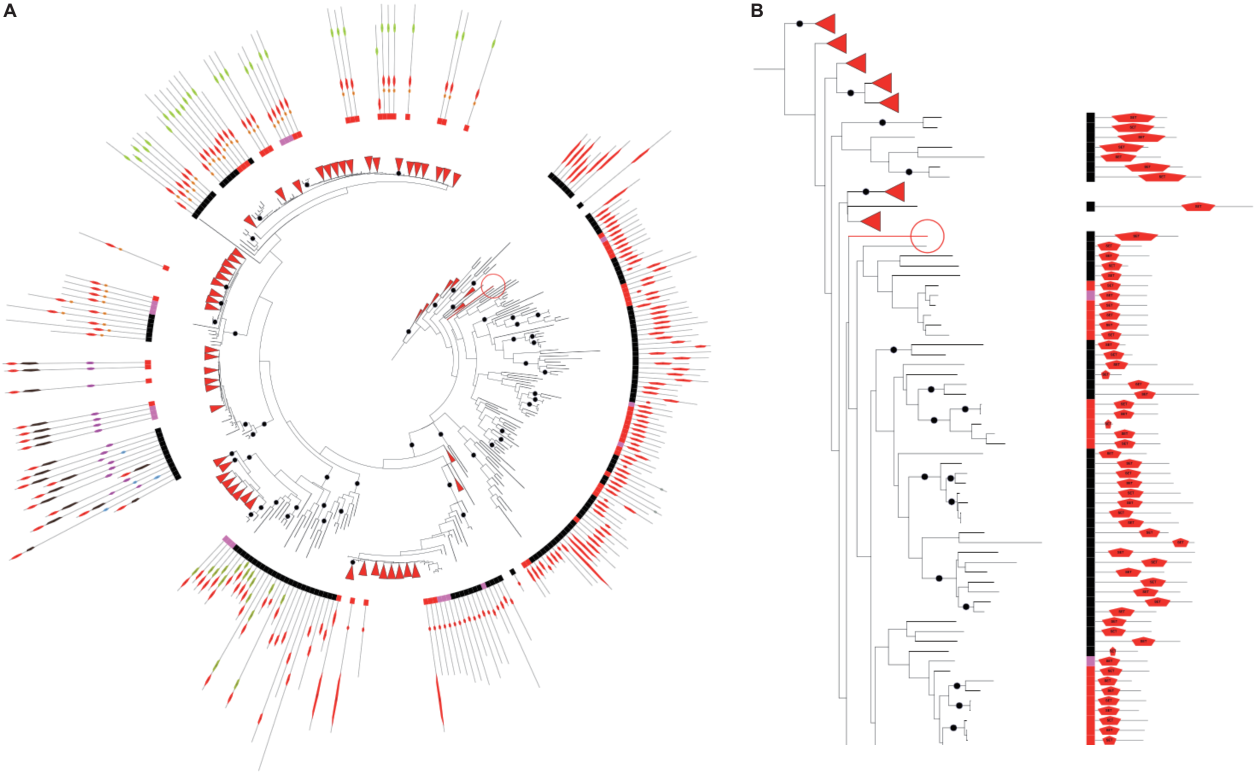


**Supplementary Figure 2 SET5 was lost in Aspergilli. A** The phylogeny of the SET domain-containing proteins in 94 Aspergilli and 15 outgroup species (Zhang X, et al., Microbial Genomics, 2022). The red circle indicates SET5 in *S. cerevisiae*. **B** Zoom-in of *S. cerevisiae* SET5 and its homologs. Black, pink, and red box indicates outgroup species, Penicillium, and Aspergilli, respectively.
