## Supplementary material for "Detection and quantification of the histone code in the fungal genus *Aspergillus*": Figure S3

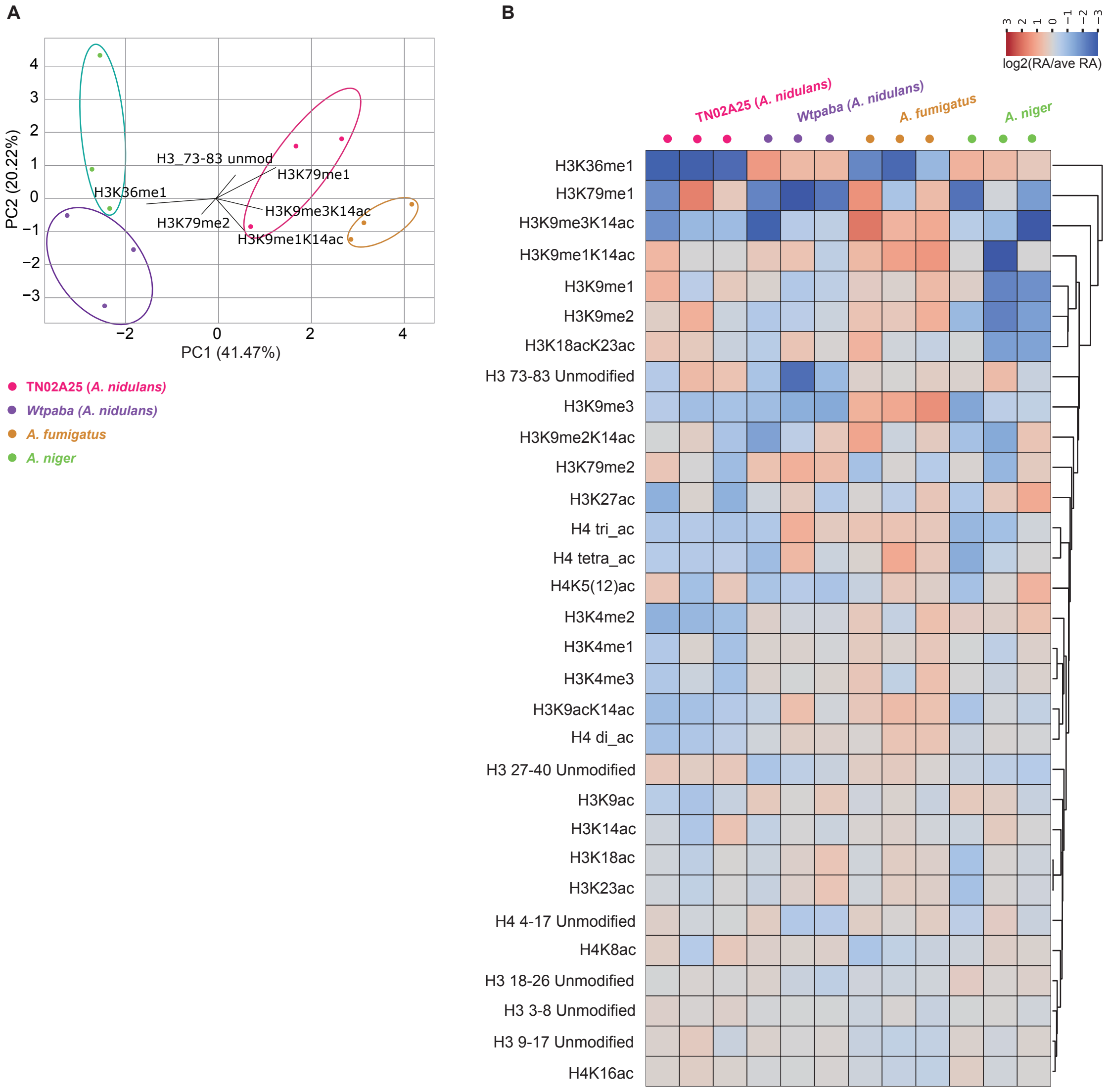


**Supplementary Figure 3 Distinct abundant patterns of several histone PTMs differ different Aspergillus species/strains. A**. Principal component analysis (PCA) according to log2(relative abundance/average of relative abundance) uncovers the separation of four strains based on their histone post translational modifications. **B**. Quantitative comparison of all histone modifications detected via mass spectrometry. The heatmap displays red or blue color boxes according to the log2(relative abundance/average of relative abundance) of different peptides in four Aspergilli. 'Di-ac' indicates H4K12acK16ac, H4K5acK12ac, H4K5acK16ac, H4K5acK8ac, or H4K8acK16ac; 'tri-ac' means H4K5acK12acK16ac, H4K5acK8ac12ac, H4K5acK8acK16ac, or H4K8acK12acK16ac; and 'tetra-ac' indicates H4K5acK8ack12acK16ac.
